# Age Associated Increase in Microglia Inflammation and Phagocytosis in the Adult Neural Stem Cell Niche

**DOI:** 10.1101/2025.10.10.681517

**Authors:** Ronald Cutler, Samuel Harrison, Natalia Sandoval-Kuhn, Stephen Tomlinson, Erzsebet Kokovay

## Abstract

Adult SVZ neurogenesis declines with age, but the niche mechanisms remain incompletely defined. Here, we show that SVZ microglia acquire activation-associated morphology with aging in male and female C57BL/6 mice. Reanalysis of published SVZ single-cell RNA-Seq data revealed that aged microglia upregulate inflammatory and lysosome/phagosome-related gene programs, including multiple phagocytosis-associated receptors and effectors. Functionally, young SVZ-derived microglia exhibit reduced basal phagocytic activity compared with non-SVZ microglia *in vitro*, whereas aged SVZ microglia show increased phagocytosis. *In vivo* SVZ wholemounts reveal increased progenitor-derived material within lysosomal compartments of microglia with age, without an accompanying increase in apoptosis. Inhibiting complement C3 activation partially restores SVZ proliferation in aged mice and coincides with a reduction of progenitor-derived material within lysosomal compartments of aged microglia. These data support a model in which microglial inflammatory signaling and engagement with neural progenitors contributes to age-related suppression of SVZ proliferation.

**Significance Statement:** Adult neural stem cells support plasticity and may aid repair after brain injury, but this capacity declines with age. We show that microglia in the aging subventricular zone acquire inflammatory and phagocytic programs, increase their lysosomal association with progenitor-derived material, and are linked to reduced progenitor proliferation. Complement inhibition partially restores proliferative output in aged mice, connecting inflammatory signaling to this niche defect. These findings identify age-dependent microglia–progenitor interactions as a mechanism that may limit neural stem cell activity in the adult brain and suggest immune pathways as possible targets for preserving or restoring neurogenic potential during aging.

## Introduction

Adult neural stem cells (NSCs) reside in two principal neurogenic niches of the rodent brain: the subgranular zone (SGZ) of the dentate gyrus and the subventricular zone (SVZ) lining the lateral ventricles. In the current study, we are focused on the NSCs in the SVZ niche, which exist in quiescent (qNSCs) and activated (aNSCs) states and generate transit amplifying cells (TACs), that expand and then produce migratory neuroblasts (Alvarez-Buylla & García-Verdugo, 2002; Doetsch et al., 1999; Lim & Alvarez-Buylla, 2016). These neuroblasts migrate from the SVZ through the RMS into the OB where they differentiate and integrate into local circuitry (Alvarez-Buylla & García-Verdugo, 2002). With aging, multiple studies report reduced SVZ neurogenesis accompanied by decreased NSC proliferation, increased NSC quiescence, and diminished NSC pool size (Ahlenius et al., 2025; Cutler & Kokovay, 2020; Enwere et al., 2004; Kalamakis et al., 2019; Maslov et al., 2004; Solano et al., 2016). These findings motivate work aimed at identifying niche-derived mechanisms that constrain activation and lineage progression in the aged SVZ.

Microglia are brain-resident macrophage-lineage cells that survey the parenchyma and shape neural homeostasis through debris clearance, phagocytosis, synapse remodeling, and secretion of signaling factors (Deczkowska et al., 2018). Fate-mapping indicates that microglia originate from early embryonic yolk sac–derived myeloid progenitors that seed the central nervous system and largely self-maintain thereafter (Ginhoux et al., 2010). Microglia constitute a substantial fraction of brain cells (commonly estimated at ∼5–15% depending on region and quantification) and display pronounced context-dependent states across development, aging, and disease (Deczkowska et al., 2018).

Several studies suggest that microglia within the SVZ are not interchangeable with microglia in non-neurogenic parenchyma. SVZ-associated microglia exhibit distinct morphological and antigenic features and have been implicated in regulating neurogenesis and progenitor dynamics (Goings et al., 2006; Marshall et al., 2014; Shigemoto-Mogami et al., 2014; Solano et al., 2016; Xavier et al., 2015). Aging is accompanied by microglial state changes in the SVZ, including increased inflammatory and lysosomal programs that precedes the decline of NSC proliferation, raising the possibility that altered microglia–NSC interactions contribute to reduced neurogenesis in the aged SVZ (Solano et al., 2016). Intriguingly, these age-related shifts in microglia towards an activated state are reminiscent of their phenotype in the developing cerebral cortex and SVZ where they have been shown to phagocytose neural precursor cells (Cunningham et al., 2013; Morrison et al., 2023). Furthermore, in a model of cortical ischemic stroke, microglia were reported to phagocytose dying neuroblasts, constraining the endogenous neurogenic response after injury (Nath et al., 2024). Despite evidence that niche inflammation can constrain activation and lineage output, it remains unclear whether aging-associated microglial activation in the SVZ is coupled to changes in microglial phagocytic behavior toward specific NSC populations, and whether inflammatory signaling and phagocytic engagement represent linked or separable mechanisms.

In this study, we test the hypothesis that aging alters SVZ microglial state and microglia–NSC interactions in ways that contribute to diminished progenitor proliferation. We found that during aging, SVZ microglia are characterized by increases in a lysosomal marker and enlarged morphologies, alongside upregulation of pro-inflammatory and phagocytic gene expression signatures, consistent with a pro-phagocytic phenotype. Aged SVZ microglia show increased phagocytosis *in vitro* and *in vivo*, where we found evidence for the engulfment of aNSCs/TACs. Finally, inhibition of complement C3 activation partially restored proliferative measures and reduced aNSC/TAC engulfment, suggesting that microglial inflammatory signaling and progenitor phagocytosis contributes to reduced NSC proliferation during aging.

## Materials and Methods

### Animals and Interventions

Male and female C57BL/6 mice were used in this study either bred at the UTHSA vivarium or acquired from the National Institute of Aging. Young mice (3 months) and aged mice (12 months) were housed in accordance with the University of Texas Health San Antonio Institutional Animal Care and Use Committee and performed in accordance with institutional and federal guidelines and allowed free access to water and standard rodent chow ad libitum.

Both male and female C57BL/6 mice received CR2-crry (10µg/1g mouse; Stephen Tomlinson Lab, Medical University of South Carolina) with saline controls via intraperitoneal injection. CR2-crry or saline was administered every other day for a total of 3 doses.

### Immunohistochemistry

The SVZ was isolated for wholemount (WM) tissue analysis; mice were anesthetized with isoflurane, a cervical dislocation was performed, followed by brain extraction. Brains were placed into cold HIB where they were halved and the septum and hippocampus were removed, and the SVZ was microdissected from the surrounding brain tissue. The SVZ WM was then fixed in fresh 4% paraformaldehyde (PFA) for 30 minutes. SVZ WMs were permeabilized in 2% PBST and blocked overnight at 4°C in 10% normal donkey serum (NDS). The SVZ WMs were then incubated at 4°C for 4 days with primary antibody in blocking solution, followed by overnight incubation at 4°C with secondary antibody made in blocking solution, and a 5-minute incubation at room temperature (RT) with DAPI (1:1000; Invitrogen, Cat. No. D1306) then mounted on a slide in ProLong Gold antifade reagent (Invitrogen, Cat. No. P36930). Primary antibodies that were used include Iba1 (1:200; Wako, Cat. No. 019-19741 and 1:200; Novus, Cat. No. NB100-1028), CD68 (1:250; BioRad, Cat. No. MCA1957), and Mash1 (1:5000; CosmoBioUSA, Cat. No. SK-T01-003) Cleaved caspase 3 (1:200 Cell Signaling Cat. No D175 5A1E). Species-specific secondary antibodies (Jackson ImmunoResearch) were used at a concentration of 1:500 in SVZ WMs.

### Immunohistochemistry Quantification

EdU was quantified in the SVZ by making a “Z Project” 2-dimensional image of the Z-stack in ImageJ on 10 random fields of view and hand counting the number of EdU+ nuclei. The number of microglia were quantified by hand counting Iba1+ cells, excluding cells whose cell body touched the image boundary on Zen Microscopy Software (Zeiss). Total CD68 area was quantified by making a “Z project” 2-dimensional image of the Z-stack, using the “Threshold” function, and then the “Analyze particle” function to give total area in µm2 that was then averaged by total number of microglia in each image. Total number of Iba1+ microglia were quantified similarly to EdU+ cells described above. Mash1 colocalization within Iba1+ microglia were quantified by hand counting in Zen Microscopy Software (Zeiss) by examining each Iba1+ microglia’s lysosome (CD68+) to determine if each Iba1+CD68+ microglia contained Mash1 colocalization. The number of Mash1+CD68+Iba1+ microglia were then averaged to total number of Iba1+ microglia in each image. There were no microglia that contained Mash1 that were not CD68+. SVZ WMs were analyzed from 15µm thick Z-stacks (15 Z-stacks per WM). Confocal images were obtained with a Zeiss LSM 710 confocal microscope at 40x objective.

### Single-cell RNA-sequencing quality control and filtering

Quality control of downloaded single-cell RNA-Seq data (see Data Availability) was first performed, which filtered cells where the number of genes or UMIs detected were greater or less than 5 median absolute deviations (MADs) or percent mitochondria reads greater or less than 3 MADs, relative to all cells in the dataset. Genes were filtered if detected in less than 10 cells out of all cells in the dataset. Cell identity labels were those assigned by the original analysis.

### Single-cell RNA-sequencing differential expression and gene set enrichment analysis

Pseudobulk samples were created by summing counts of each identified cell type for each biological replicate. The DESeq2 package (Love et al., 2014) was then used to perform differential expression analysis to compare old vs. young groups. Following this the ClusterProfiler package (Yu et al., 2012) was then used to perform gene set enrichment analysis using the gene ontology biological process database or custom annotations.

### Microglia phagocytosis gene set analysis

Genes associated with known phagocytosis receptors and/or pathways active in microglia were manually curated by searching the Molecular Signatures Database (MsigDB) and the literature (i.e. Complement, Toll-like receptor, TAM receptor, Trem2 receptor, FC gamma, Pr2y6 receptor, Scavenger receptor, Mannose receptor, LRP1 receptor, Galectin-3 adaptor protein, Microglia Activation, Microglia Proliferation, Disease-Associated Microglia (DAM), and Microglia Development). In addition, gene sets related to screens of phagocytosis regulators was obtained from the literature (see Data Availability). These gene sets were then overlapped with the genes that were found to be significantly differentially expressed (adjusted p-value < 0.05) in the microglia old vs. young comparison.

### Neural stem cell gene set analysis

Genes associated with pathways known to be expressed in cells that are targeted for phagocytosis were manually curated by searching the Molecular Signatures Database (MsigDB) and the literature (i.e. Apoptosis, Opsonization, Antigen Processing and Presentation, “eat-me,” “find-me,” and “don’t eat me”). Because of the lack of genes which were significantly differentially expressed in neural stem cells (i.e. Astrocytes/qNSCs, aNSCs/NPCs and Neuroblasts) in the old vs. young comparison, the ClusterProfiler package was then used to perform gene set enrichment analysis using the manually curated gene sets. The Log_2_ fold change of genes within the gene sets found to be significantly enriched in age-related genes expression of neural stem cells.

### Microglia Isolation and Cell Culture

SVZ microglia were derived by mixed-glial expansion whereas whole brain microglia were MACS-isolated. C57BL/6 mice were injected intraperitoneally with pentobarbital (25µL/mouse) then perfused with cold 1x PBS. SVZ WMs or the remainder of the whole brain excluding the SVZ and cerebellum, hereby referred to as whole brain (WB), were isolated following perfusion. The tissues were minced and dissociated in 50U papain in DMEM/F12 at 37°C for 45 minutes and manually triturated and spun 3x in DMEM. SVZ and WB homogenate were gently pressed through a 70µm cell strainer to remove debris. For mixed glial culture, SVZ homogenate were then plated in a T25 flask coated with poly-L-ornithine (PLO) in neural growth media (DMEM/F12, 5% fetal bovine serum, 1x N2, 10ng/mL EGF, and 10ng/mL bFGF) until a confluent cell monolayer formed. The culture was then fed microglia proliferation media (DMEM/F12, 10% fetal bovine serum, 1x N2, and 20 ng/mL GM-CSF) until microglia could be seen atop the monolayer. Once microglia were present, they were shaken off from the cell monolayer and the SVZ microglia could be plated for further experimentation.

For microglia isolated by MACS, following the cell strainer step, WB homogenate was resuspended in HIB and centrifugated at 400G with no brake in 22% Percoll (Sigma-aldrich, Cat. No. P4937) solution to remove myelin from the samples. The WB homogenate was then resuspended in 0.5% bovine serum albumin and incubated with 10µL CD11b Microbeads (cat# 130-093-636) per 107 cells. The homogenate was then placed in an LS column (MiltenyiBiotec, Cat. No. 130-042-401) in a QuadroMACS™ separator (MiltenyiBiotec, Cat. No. 130-090-976) to allow for MACS sorting of CD11b + microglia following manufacturer protocols (MiltenyiBiotec). WB microglia were plated in a 6-well plate coated with PLO/laminin (10µg/mL; Sigma-aldrich, Cat. No. L2020) at 250k cells/well in microglia expansion media (DMEM/F12, 100 µg/mL PenStrep, 5 µg/mL n-acetyl-L-cysteine, 5 µg/mL Insulin, 100 µg/mL apo-transferrin, 100 µg/mL sodium selenite, 1 µg/mL heparin sulfate, 10% fetal bovine serum, 1.5 µg/mL cholesterol, 40 ng/mL M-CSF, 20 ng/mL GM-CSF, and 2 ng/mL TGF-β) adapted from [Bohlen et al., 2017].

### Microglia *in vitro* Phagocytosis Assay

The phagocytosis assay was performed using live cell imaging by Incucyte S3 (Essen BioScience) for WB or SVZ microglia phagocytosis from 3- or 12-month mice. WB microglia were passaged from a 6-well plate with Trypsin (Fisher, Cat. No. 25-200-072) following MACS isolation and SVZ microglia were collected from mixed glial culture and microglia were plated in a 96 well plate coated with PLO/laminin at 20k cells/well in microglia expansion media for 3 days. After 3 days, SVZ and WB microglia were moved to serum free media (DMEM/F12, 100 µg/mL PenStrep, 5 µg/mL n-acetyl-L-cysteine, 5 µg/mL Insulin, 100 µg/mL apo-transferrin, 100 µg/mL sodium selenite, 1 µg/mL heparin sulfate, 1.5 µg/mL cholesterol, 40 ng/mL M-CSF, and 2 ng/mL TGF-β) adapted from [Bohlen et al., 2017] for 1 week. All microglia were then incubated with 5 µg/mL Isolectin IB4 Alexa Fluor™ 488 Conjugate (Invitrogen, Cat. No. I21411) for 60 minutes at 37°C. SVZ or WB microglia of both ages were treated with Zymosan S. cerevisiae BioParticles™ (10 mg/mL; Invitrogen, Cat. No. Z2849) conjugated pHrodo™ Red SE (Molecular Probes™, Cat. No. P36600) following manufacturer protocols (ThermoFisher) and adapted from [Haney et al., 2018; Pluvinage et al., 2019], both Zymosan-pHrodo and Cytochalasin D (10 mM/µL Sigma-aldrich, Cat. No. C8273) to inhibit phagocytosis, or isolectin only as a negative control. The microglia were then incubated in the Incucyte S3 with fluorescent images captured every hour over the course of 48 hours for live cell imaging of microglia phagocytosis. The images were analyzed in the IncuCyte software (Essen BioScience) quantifying colocalization of the green fluorescence of Isolectin IB4 labelled microglia and the red fluorescence of engulfed Zymosan-pHrodo within the microglia lysosome demonstrating microglia phagocytosis. Total pHrodo area / number of microglia was calculated for each well and condition yielding amount of in vitro phagocytosis occurring at each time point.

### RNA Extraction and qPCR

RNA was extracted from freshly isolated microglia following magnetic-activated cell sorting (MACS) isolation, using a TRIzol-chloroform method (Rio et al., 2010) where cells were lysed in TRIzol (Invitrogen, Cat. No. 15596018), aqueous layer separated with chloroform, RNA was precipitated with isopropanol, washed in ethanol, dried, then resuspended in RNase-free water. cDNA was synthesized using High Capacity cDNA Reverse Transcription Kit (Applied Biosystems, Cat. No. 4368814). Quantitative polymerase chain reaction (qPCR) was performed using a C1000 Touch Thermal Cycler (BioRad) with iTaq™ Universal SYBR® Green Supermix (BioRad, Cat. No. 1725121). Reactions were performed in doublet with glyceraldehyde-3-phosphate dehydrogenase (GAPDH) used as an endogenous control. Negative control reactions were performed using the same conditions without cDNA. Quantification was analyzed using the ΔΔCt method as described [(Livak & Schmittgen, 2001)]. Primers were used for the following transcripts: C1QA, P2Y12, TREM2, LPAR, C3, CR1, ITGAM, ITGAX, GAPDH.

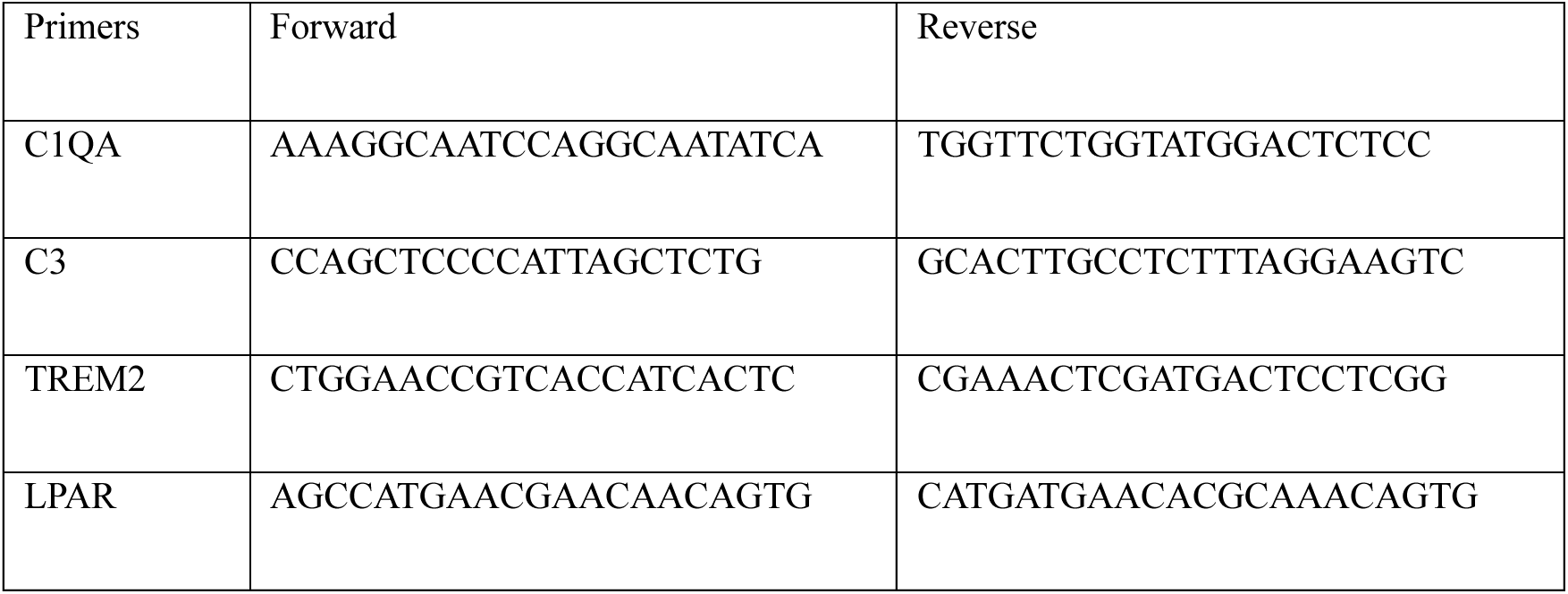
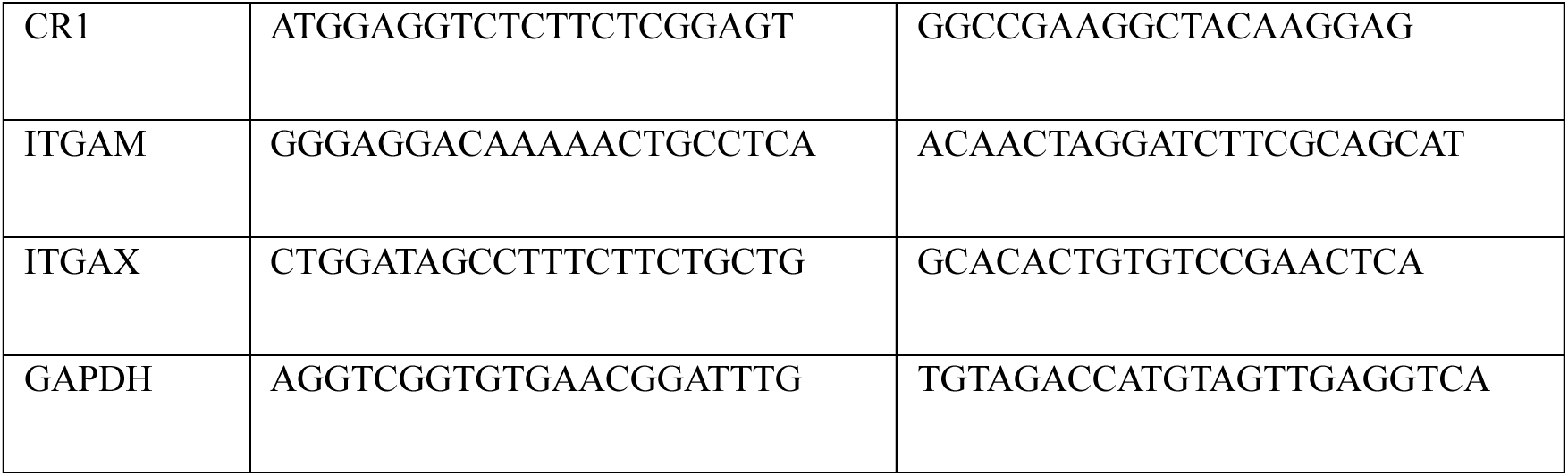

### EdU Labeling and Staining

Mice were injected intraperitoneally (50µg EdU/1g mouse) 2 hours before sacrifice. Their brains were removed and SVZ was isolated and fixed for 30 minutes in 4% PFA. EdU staining was performed following manufacturer protocols (Fisher, Click-it EdU Imaging Kit, Cat. No. C10337).

### Experimental Design and Statistical Analysis

Student’s t-tests and two-way ANOVA were used to analyze the data on GraphPad Prism 8 Software. All data are expressed as mean ± standard error of the mean.

### Data Accessibility

Normalized single-cell RNA-sequencing data and cell type annotations of the SVZ from young (3 months old) and old (28–29 months old) male mice (n = 3) was downloaded from Supplementary Table 2 of the Dulken et al., 2019 study. Results of screen for phagocytosis regulator genes were downloaded from Supplementary Table 5 of the Haney et al., 2019 study and Supplementary Table 2 of the Pluvinage et al., 2019 study. Marker genes of major cell types in the SVZ of young (2-4 months old) male and female mice were downloaded from Table S1 of the Zywitza et al., 2018 study.

### Code Accessibility

Code to reproduce single-cell RNA-sequencing data analyses is available at the following repository: https://github.com/cutleraging/nsc-aging

## Results

### Age-related changes in SVZ microglia morphology

We have previously reported that murine SVZ neurogenesis is reduced by 12 months of age (Solano et al., 2016). To assess how SVZ microglia change during aging, we performed immunohistochemistry on SVZ wholemount preparations (to preserve 3D niche cytoarchitecture) from young (3 month) and aged (12 month) mice, staining for the microglial marker Iba1 and the lysosomal marker CD68. In young mice, Iba1+ microglia exhibited a ramified morphology throughout the SVZ with relatively low CD68 signal within Iba1+ cells (Fig. 1A-B). In aged mice, Iba1+ microglia displayed morphology consistent with activation, including fewer processes and enlarged somata, together with increased CD68 signal within Iba1+ cells (Fig. 1C-D). Because microglia can adopt a lysosome-rich, phagocytic phenotype during neuronal development that is associated with engulfment of neural precursor cells, we hypothesized that age-associated microglial activation in the SVZ is accompanied by increased phagocytic activity toward SVZ progenitors (Cunningham et al., 2013).

**Figure 1.**
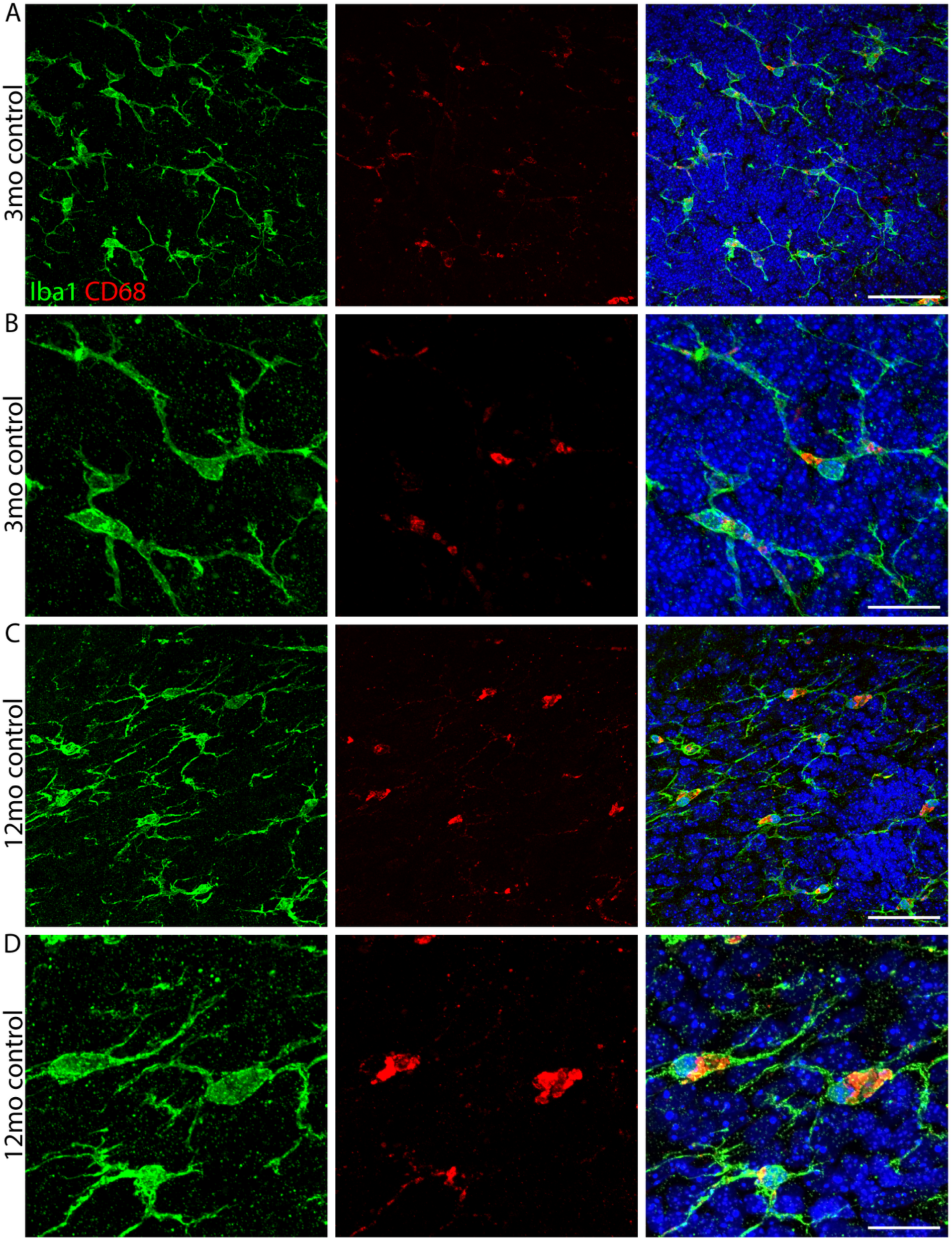
Age-Associated SVZ Microglia Morphological Changes and Increased Activation. Representative confocal projection images of SVZ microglia stained with Iba1 (green), lysosomes stained with CD68 (red), and nuclei stained with DAPI (blue). **(A-B)** Young (3 month) SVZ microglia imaged at 40X magnification (A) or 100X magnification (B). **(C-D)** Aged (12 month) SVZ microglia imaged at 40X magnification (C) or 100X magnification (D). Scale bars in (A) and (C) = 20µm, scale bars in (B) and (D) = 50µm

### Gene expression in the aging SVZ suggests increased microglial phagocytic programs

To assess age-related transcriptional changes in SVZ microglia and NSC populations relevant to inflammation and phagocytosis, we reanalyzed a published single-cell RNA-Seq dataset comprising of 14,685 cells from the SVZ of young (3 month) and old (28/29 month) male mice (Dulken et al., 2019). We generated pseudobulk profiles for each cell type and biological replicate, performed differential gene expression analysis (old vs. young), followed with gene set enrichment analysis (GSEA) (Supplementary Figs. 1A-B).

In microglia, we identified 254 genes significantly upregulated and 54 genes significantly downregulated with age (adjusted p-value < 0.05; Supplementary Fig. 1C; Supplementary Table 1). GSEA of genes upregulated with aging revealed enrichment of processes related to translation, secretion of inflammatory cytokines, and lipid metabolism, consistent with prior analyses (Dulken et al., 2019; Yen & Yu, 2024). In contrast, downregulated genes were enriched for cytoskeleton organization terms, which has not been previously reported and may suggest dysfunction in motility or morphology-related programs (Supplementary Fig. 1D).

Although canonical “phagocytosis” terms were not significantly enriched by GSEA (Supplementary Fig. 1D), several phagocytosis-associated receptors, including *Axl* and *Tlr2*, were among the most strongly upregulated transcripts with age (Supplementary Fig. 1C). We therefore curated comprehensive gene sets representing multiple components of phagocytosis from MSigDB (Liberzon et al., 2024) and intersected these with age-related differentially expressed genes (ARDEGs; adjusted p-value < 0.05; Supplementary Table 2). 19 phagocytosis-related genes overlapped with the microglia ARDEG set, of which 18 were upregulated in aged microglia (Fig. 2A; Supplementary Table 3). Notably, 8 of these genes (*Cyba, Atp6v0e, Rab7b, Cybb, Vim, B2m, Tlr2,* and *Hvcn1*) are linked to phagosome formation/maturation and vesicular acidification. We further expanded our phagocytosis gene list to include genes identified in prior functional screens (Haney et al., 2018; Pluvinage et al., 2019), and found that all intersecting genes (9/9) were upregulated in aged microglia (Supplementary Figs. 2A-B). Finally, we found that the broad upregulation of phagocytosis-related gene expression was also accompanied by an increase in the expression of genes related to microglia activation (Supplementary Figs. 2C-D). While these transcriptional shifts are consistent with increased lysosomal/phagocytic programs, we note that elevated expression of such genes could also reflect compensatory responses to altered phagocytic demand or efficiency rather than a direct readout of increased engulfment per se. For instance, the expression of the receptor essential for microglial function, *Csf1r*, was downregulated in aged microglia while the ligand *Csf1* showed the opposite trend and thus may be a compensatory response to chronic microglia activation (Supplementary Fig. 2C) (Hu et al., 2024). Together, these findings suggest that aging is accompanied by transcriptional reprogramming of SVZ microglia toward states featuring inflammatory signaling and lysosome/phagosome-related programs, with partial resemblance to developmental and phagocytic microglial states (Cunningham et al., 2013; Izgi et al., 2022).

**Figure 2.**
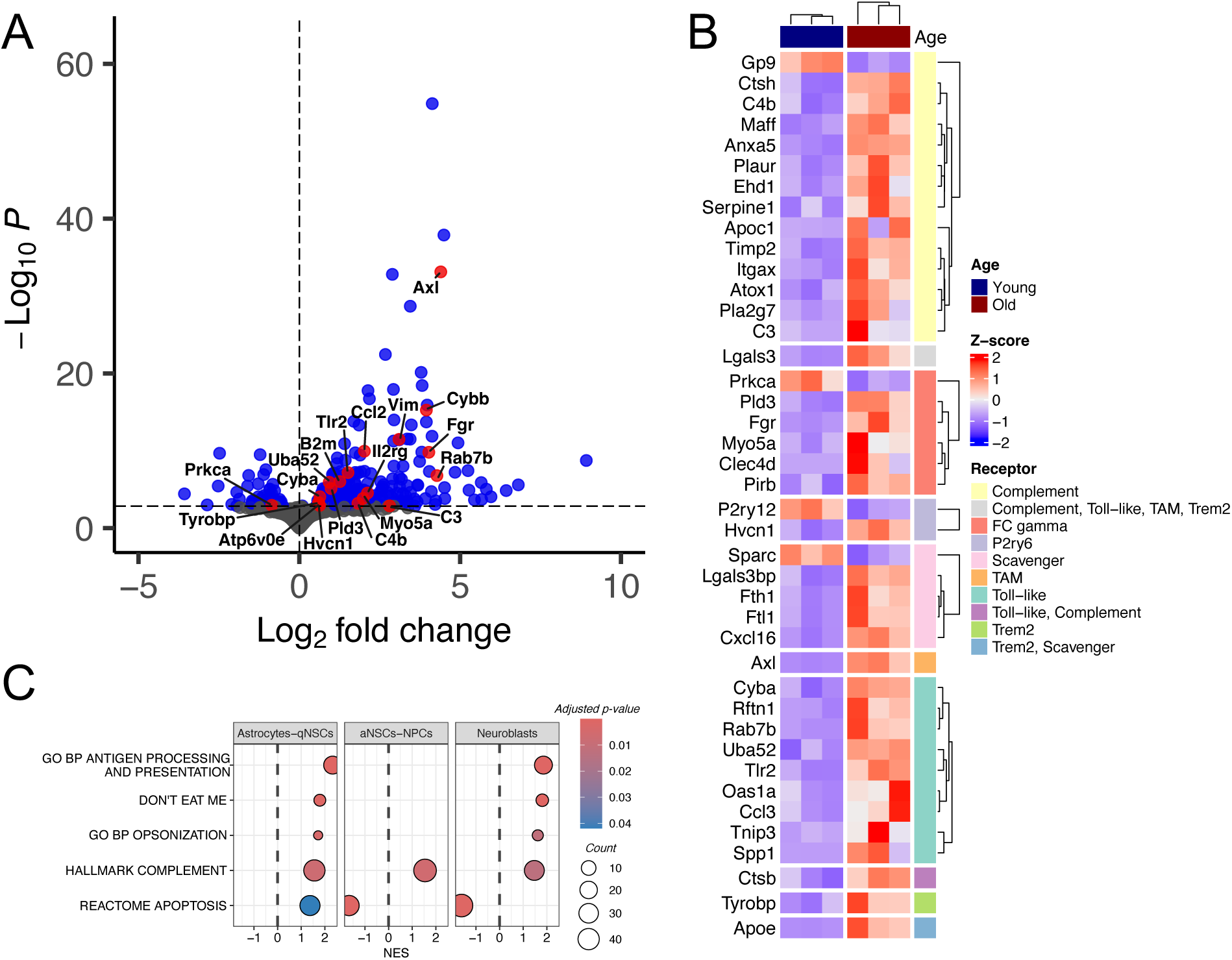
Gene expression in the aging SVZ suggests increased microglial phagocytic programs. **(A)** Pseudobulk level analysis of differential gene expression changes during aging of young (3 month) and aged (28–29 month) microglia in the SVZ. Red points indicate genes that overlapped with the phagocytosis-related gene sets (Supplementary table 2) which have absolute log2 fold change > 1 and adjusted p-value < 0.05. Statistics: n=3 mice/group, DGE results obtained using Wald test followed by Benjamini-Hochberg correction for multiple hypothesis testing. **(B)** Heatmap of age-related DEGs (adjusted p-value < 0.05) in aging SVZ microglia (obtained from analysis in (A)) that overlapped with phagocytosis-related gene sets (Supplementary table 2) Statistics: n=3 mice/group. **(C)** Gene set enrichment analysis of age-related gene expression changes in neural progenitor cells using apoptosis, “find-me,” “eat-me,” and “don’t eat-me” signal gene sets (Supplementary table 2). Statistics: n=3 mice/group, GSEA p-values obtained using a permutation test followed by Benjamini-Hochberg correction for multiple hypothesis testing.

We next asked whether age-associated changes were concentrated within particular receptor-mediated phagocytosis pathways. Using a curated list for the major phagocytosis pathways, we observed upregulation of the receptors in the Complement, TAM, and Toll-like pathways (Fig. 2B; Supplementary Table 3). In contrast, for the FC gamma, Scavenger, P2ry6, and Trem2 receptor pathways, multiple pathway components were consistently upregulated without a corresponding increase in the core receptor transcript (Deczkowska et al., 2018). Notably, while Trem2 itself was not increased, aged microglia displayed expression patterns consistent with a disease-associated microglia (DAM) stage 2 signature that has been reported to be Trem2-dependent in other contexts (Supplementary Fig. 2E). Overall, these results indicate broad multi-pathway upregulation of phagocytosis-related gene expression during aging and suggest that aged SVZ microglia may adopt a unique Trem2-independent DAM-like state.

We then analyzed NSC populations for age-associated expression changes in signals relevant to target-cell recognition and engulfment, including apoptosis, “find-me,” “eat-me,” and “don’t eat-me” programs. Compared to microglia, there were very few ARDEGs: 13 genes were significantly upregulated in aged astrocyte-qNSC as compared to young (adjusted p-value < 0.05), whereas no genes met this threshold for aged aNSC, TACs or neuroblasts (Supplementary Figs. 1E-J). Given this limited power for threshold-based overlap analyses, we used GSEA with curated gene sets for apoptosis and phagocytosis-related signaling (Supplementary Table 2).

Using this approach, we detected age-associated upregulation of an apoptosis gene set in astrocyte-qNSC cells (adjusted p-value < 0.05), whereas aNSC, TACs and neuroblasts showed the opposite trend (Fig. 2C; Supplementary Fig. 3A). We did not detect significant enrichment for curated “find-me” or generic “eat-me” gene sets. However, we observed age-associated upregulation of complement system and opsonization gene sets (often considered “eat-me”–adjacent signals) in astrocyte-qNSCs and neuroblasts, but not in aNSCs or TACs (Fig. 2C; Supplementary Fig. 3B,C) (Li, 2012). Intriguingly, astrocyte-qNSCs and neuroblasts also showed increased expression of a curated “don’t eat-me” gene set (Fig. 2C; Supplementary Fig. 3D). This set includes genes related to MHC class I antigen presentation such as *H2-K1* and *H2-D1*, which in some contexts can oppose macrophage phagocytosis, particularly when expressed with *B2m* (Supplementary Fig. 3D) (Khalaji et al., 2023). Consistent with a potential receptor–ligand axis, *Lilrb4a*, an inhibitory receptor for MHC class I, was among the age-upregulated microglial ARDEGs (Supplementary Fig. 1C). An antigen processing/presentation gene set showed a similar pattern to the “don’t eat-me” set, further supporting increased MHC class I–associated programs in aged astrocyte-qNSCs and neuroblasts, but not aNSC-NPCs (Supplementary Fig. 1E).

In summary, transcriptomic changes in SVZ microglia with age are consistent with increased inflammatory and lysosome/phagosome-related programs across multiple receptor pathways. In aged NSC populations, astrocyte-qNSCs and neuroblasts showed increased complement pathway components (“eat-me” signals) together with increased expression of putative “don’t eat-me” anti-phagocytic signals, while aNSC-NPCs did not show comparable upregulation of those signals. These patterns are consistent with a model in which age-associated microglial phagocytic programs may preferentially engage aNSC/TACs.

### SVZ microglia show an age-associated increase in phagocytic activity toward SVZ progenitors

To assess SVZ microglia phagocytic activity, we microdissected the SVZ and isolated microglia either from the SVZ or from the remainder of the brain (excluding the SVZ) from young (3 month) and aged (12 month) mice. Isolated microglia were expanded *in vitro*, and phagocytosis was quantified by live imaging of engulfment of pHrodo-conjugated yeast particles (pH-sensitive reporter that fluoresces upon acidification in phagolysosomes) (Kapellos et al., 2016). In young animals, SVZ-derived microglia displayed lower phagocytic activity than microglia isolated from non-SVZ brain regions under our assay conditions, consistent with niche-associated specialization toward reduced basal phagocytosis (Fig. 3A) (Xavier et al., 2015). In aged animals, SVZ-derived microglia showed a significant increase in phagocytic activity relative to young SVZ microglia, approaching the levels observed in whole-brain microglia.

**Figure 3.**
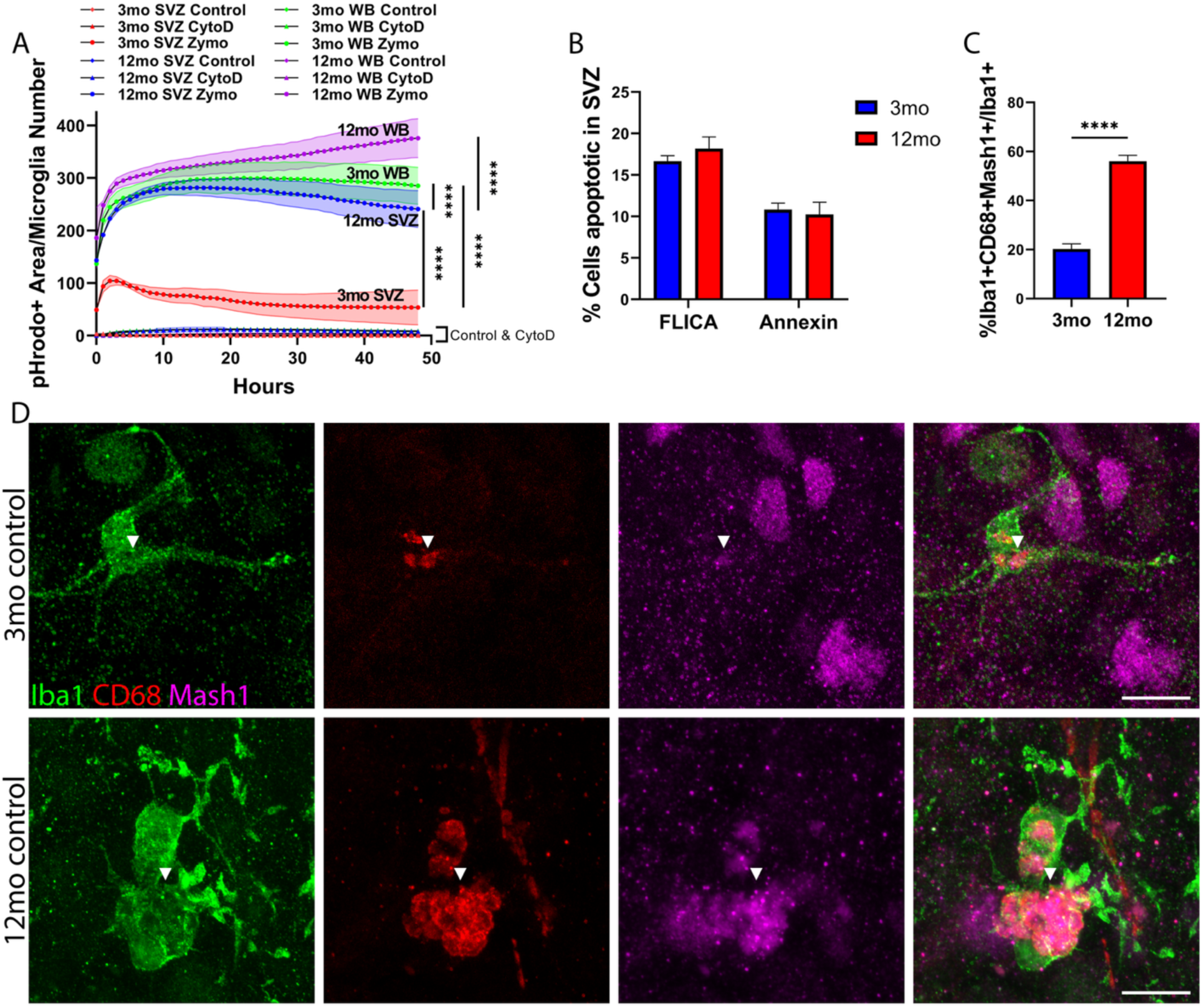
SVZ microglia show an age-associated increase in phagocytic activity toward SVZ progenitors. **(A)** Time-lapse *in vitro* phagocytosis assay of young (3 months) or aged (12 months) SVZ or whole brain (WB) microglia labeled with isolectin and incubated with Zymosan-pHrodo (Zymo), CytochalasinD and Zymosan-pHrodo (CytoD, negative control), or isolectin only (Control). Phagocytosis is quantified as pHrodo+ area indicated by red fluorescence / microglia number. Statistics: n=3 mice/group, ≥ 1,000 microglia quantified per mouse. (B) Quantification of apoptosis in SVZ whole mounts. Statistics: n=6 mice/group, p-value estimated using two-way ANOVA. (C) Quantification of *in vivo* phagocytosis of 3- or 12-month SVZ microglia measured as Iba1+CD68+Mash1+ microglia / total Iba1+ microglia (Methods). *n*=11 mice/group, ≥100 microglia quantified per mouse. (D) Representative confocal projection images of young or aged SVZ microglia stained with Iba1 (green), microglia lysosomes stained with CD68 (red), and TAC transcription factor stained with Mash1 (magenta). Scale bar: 10µm, p-value estimated using two-way ANOVA. P-value legend: ns: no symbol; ****: p ≤ 0.0001.

We next sought *in vivo* evidence that the age-associated increase in SVZ microglial phenotypes reflects increased phagocytic engagement with SVZ progenitors. As an initial test, we quantified apoptosis in the SVZ by immunohistochemistry for cleaved caspase-3 on coronal sections from young (3 month) and aged (12 month) mice and did not detect a significant age-associated increase in apoptotic cells in the SVZ (Fig. 3B). Next, we sought to validate the findings of an age-related increase in SVZ microglia phagocytosis activity in our *in vitro* system by examining SVZ microglia phagocytosis *in vivo*. We assessed microglial phagolysosomal content directly by staining SVZ wholemounts for Iba1 (microglia), CD68 (lysosomes), and Mash1 (aNSC/TAC specific transcription factor) (Kim et al., 2011; Lim & Alvarez-Buylla, 2016). In both young and aged SVZ, we observed Mash1 signal within CD68+ compartments inside Iba1+ microglia (Fig. 3D). Quantification indicated that ∼20% of SVZ microglia contained Mash1+ material within CD68+ compartments in young mice, and this fraction increased over 2-fold at 12 months of age (Fig. 3C). Together, these in vitro and in vivo data support an age-associated increase in SVZ microglial phagocytic activity and increased microglial phagolysosomal association with progenitor-derived material in the aged SVZ.

### Complement inhibition partially rescues SVZ proliferation in aged mice

To explore possible mechanisms that might contribute to the age-associated increase in SVZ microglial phagocytic engagement of aNSC/TAC populations in the SVZ, we examined expression of candidate pro-inflammatory factors in SVZ microglia guided by our transcriptomic reanalysis (Fig. 2A). We reasoned that pro-inflammatory genes that were low or absent in the young SVZ microglia relative to non-SVZ microglia that are increased with age in SVZ microglia would be good candidates based on our *in vitro* phagocytosis results (Fig. 3A). Among the candidates, *C3* (Complement component 3) showed higher expression in non-SVZ microglia relative to young SVZ microglia and was strongly upregulated with age in SVZ microglia (Supplementary Fig. 4). In addition, transcripts encoding *C3* receptors *Itgam* and *Itgax* were increased in aged SVZ microglia. Based on these observations, we tested whether inhibiting *C3* activation would alter SVZ proliferation and/or microglial phagocytic readouts *in vivo*.

To inhibit C3 cleavage and deposition, we used CR2-Crry, a targeted complement inhibitor that crosses the blood brain barrier and localizes to sites of complement activation via CR2 to then block C3 activation through Crry (Atkinson et al., 2005; Song et al., 2003). Young (3 month) and aged (12 month) mice received CR2-Crry or saline every other day for three total doses, followed by a 2-hour EdU pulse prior to sacrifice to label proliferating cells (Fig. 4A). SVZ wholemounts were collected and processed for immunohistochemistry as described above (Methods).

**Figure 4.**
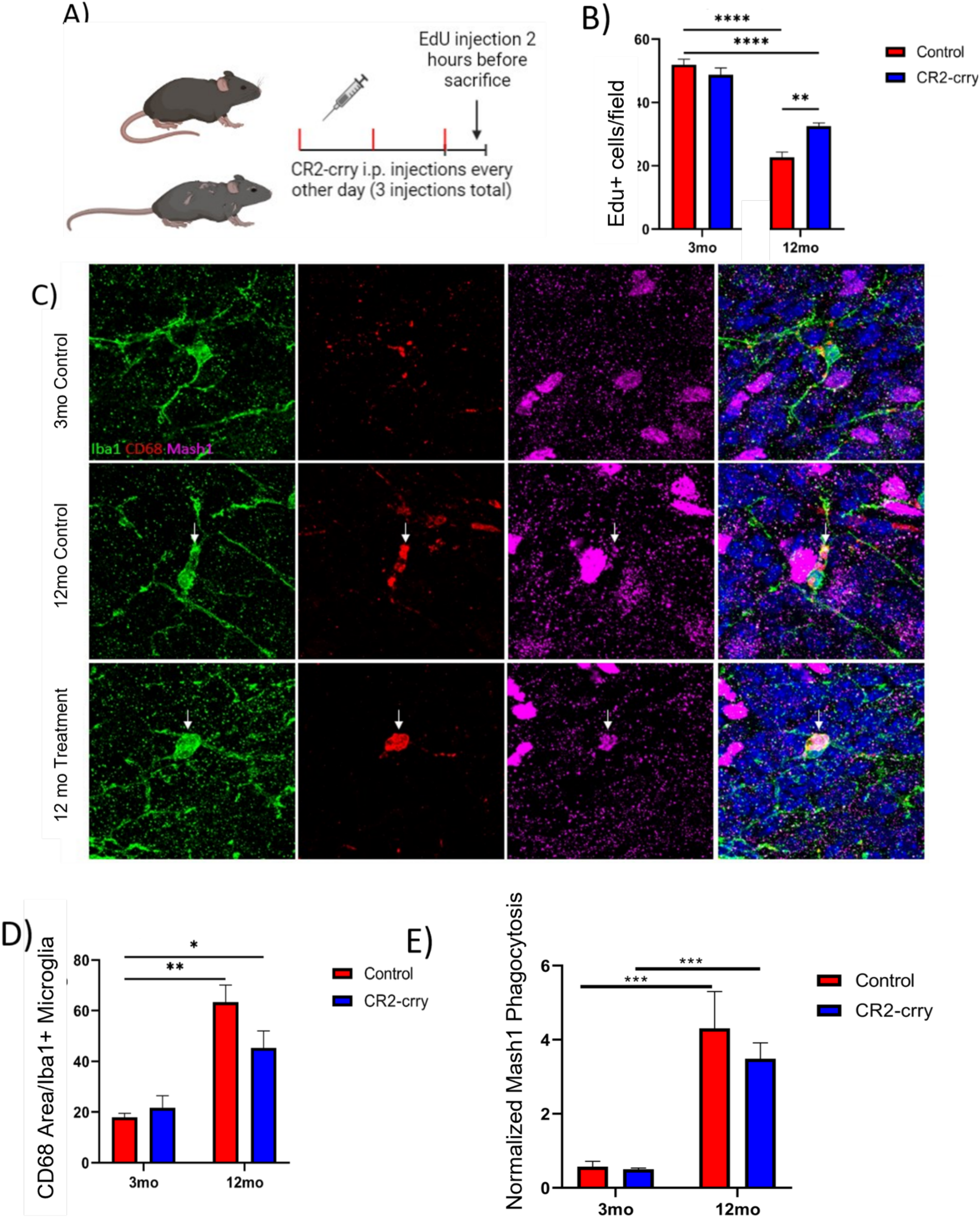
Complement inhibition partially rescues SVZ proliferation in aged mice. **(A)** Graphic representation of CR2-crry or saline administration to young (3 month) or aged (12 month) mice. **(B)** Quantification of proliferating EdU-positive cells in the SVZ of young or aged control or CR2-crry treated mice. Statistics: *n*=4 mice/group, ≥ 100 image fields quantified per mouse, p-values estimated using two-way ANOVA. **(C)** Confocal projection images of young control, aged control, aged CR2-crry treated SVZ microglia stained with Iba1 (green), microglia lysosomes stained with CD68 (red), and aNSC/TAC transcription factor stained with Mash1 (magenta). Arrowheads in aged control and aged CR2-crry treated indicate colocalization of Mash1+ signal within microglia lysosomes (Iba1+CD68+). Scale bar: 10µm. **(D)** Quantification of microglia activation measured by CD68 area / Iba1+ cells. Statistics: *n*=4 mice/group, ≥ 1,000 microglia quantified per mouse, p-values estimated using two-way ANOVA. **(E)** Quantification of Mash1 containing microglia (Iba1+CD68+Mash1+ / Iba1+ cells / Mash1+ cells) from young or aged control or CR2-crry treated mice. Statistics: Same as in (D). P-value legend: ns: no symbol; *: p ≤ 0.05, **: p ≤ 0.01, ***: p ≤ 0.001, ****: p ≤ 0.0001.

In young mice, CR2-Crry treatment did not significantly affect SVZ proliferation as indicated by EdU+ cells, microglial CD68 area, or the fraction of microglia containing Mash1+ material within CD68+ compartments (Fig. 4C). As expected, saline-treated aged mice showed reduced SVZ proliferation relative to saline-treated young mice (Fig. 4B). In aged mice, CR2-Crry significantly increased SVZ proliferation and Mash1+ cells compared with age-matched saline controls as indicated by EdU+ cells (Fig. 4B; Supplementary Fig. 5). CD68 area per Iba1+ microglia was increased in aged saline-treated mice relative to young controls, while CR2-Crry produced a non-significant reduction in this measure in aged mice (Fig. 4D). Similarly, the microglia containing Mash1+ material within CD68+ compartments increased with age, and CR2-Crry treatment resulted in a non-significant reduction relative to aged saline controls (Fig. 4E). Together, these data indicate that complement inhibition via CR2-Crry can partially restore proliferative output in the aged SVZ.

## Discussion

Adult SVZ NSCs generate new neurons that migrate to the OB and contribute to olfactory learning and discrimination throughout life (Lledo & Valley, 2016). In addition, following brain injury SVZ neural progenitors and neuroblast undergo a proliferative burst and then migrate to damaged areas where they give rise to a limited number of neurons, astrocytes and oligodendrocytes and can also secrete pro-reparative factors (Liang et al., 2019; Robin et al., 2006; Yamashita et al., 2006). The blockade of this response results in larger lesions and worsens behavioral deficits (Ohab et al., 2006). Together, these observations support the SVZ as a candidate target for therapies aimed at enhancing endogenous repair. However, SVZ neurogenesis declines drastically in aged rodents, with the most-rapid reductions between midlife (9-12 months) and persists for the duration of aging (Luo et al., 2006). This decline coincides with increasing risk of brain injury with age, motivating a need to identify niche mechanisms that constrain SVZ stem/progenitor function. Our data support a model in which aging is accompanied by increased microglial inflammatory and phagocytic programs within the SVZ niche, together contributing to reduced stem/progenitor output during aging.

During embryonic neurogenesis, SVZ-associated microglia display amoeboid morphology and can regulate precursor pool size through phagocytosis of non-apoptotic neuroblasts (Cunningham et al., 2013; Morrison et al., 2023). In the adult dentate gyrus, another neurogenic niche, microglia phagocytose apoptotic newborn neurons as part of routine homeostasis, whereas apoptosis within the adult SVZ is generally reported to be low (Luo et al., 2006). In the OB, where many SVZ-derived neurons fail to integrate, phagocytic clearance is more prominent (Xavier et al., 2015). In our study, we did not detect an age-associated increase in apoptotic signal in the SVZ, arguing against a simple model in which increased microglial lysosomal content in aging is driven primarily by increased clearance of apoptotic SVZ progenitors. Instead, our *in vivo* colocalization and *in vitro* phagocytosis assays are consistent with increased microglial phagocytic programs and increased microglial association with Mash1+ progenitor-derived material in the aged niche, although we emphasize that colocalization-based metrics are proxies and do not, on their own, establish engulfment kinetics or target selectivity.

Reanalysis of published single-cell transcriptomes from young and old SVZ provided an independent line of evidence consistent with increased inflammatory and phagocytosis-related programs in aged microglia. In pseudobulk AGDGE and gene-set analyses, aged microglia upregulated inflammatory signaling and lysosome/phagosome-associated genes, including multiple receptors and effectors linked to phagocytic function (e.g., *Axl*, *Tlr2*, *Lgal3*, and *C3*), and showed broader increases across several phagocytosis pathway components (e.g. Complement, FC gamma, and Toll-like). In contrast, neural stem/progenitor populations exhibited comparatively few significant age-related DE genes, motivating gene-set–level testing rather than threshold overlaps. Those analyses suggested that astrocyte-qNSCs and neuroblasts increase complement/opsonization-related expression (i.e. “eat-me” programs) while also increasing expression of putative anti-phagocytic programs, including MHC class I/antigen presentation–associated genes (i.e. “don’t eat-me” programs). aNSCs-NPCs did not show the same pattern of increased “don’t eat-me” programs, which is consistent with, but does not prove, a model in which aNSCs/ TACs may be relatively vulnerable to microglial phagocytic engagement in the aged niche. Importantly, transcriptomic signatures cannot distinguish increased phagocytic capacity from compensatory responses to altered substrate availability, nor can they assign directionality without functional validation. The strength of the scRNA-seq analysis is therefore as mechanistic context that complements the *in vivo* and *in vitro* assays.

Microglia are distributed throughout the brain, but exhibit region-dependent heterogeneity in density, morphology, transcriptional state, and receptor expression shaped by local microenvironments (Grabert et al., 2016; Tan et al., 2020). SVZ microglia are reported to be phenotypically distinct from non-germinal regions, supporting the idea of niche specialization (Goings et al., 2006; Marshall et al., 2014; Shigemoto-Mogami et al., 2014; Solano et al., 2016; Xavier et al., 2015). Riberio Xavier et al. showed that SVZ microglia in young adults have distinct morphology and receptor profiles, are required for neuroblast survival and migration, and display comparatively low phagocytic activity in the young SVZ niche, potentially linked to reduced expression of specific purinergic receptors (Xavier et al., 2015). Our *in vitro* assay is consistent with this niche-specific baseline state. SVZ-derived microglia from young mice exhibited lower phagocytic activity than microglia isolated from non-SVZ brain regions. With aging, we observe a shift toward increased CD68+ lysosomal features and increased phagocytic readouts, consistent with remodeling of this niche-specialized state.

To probe whether inflammatory signaling contributes causally to SVZ aging phenotypes, we used CR2-Crry, a targeted inhibitor of complement C3 activation and deposition (Atkinson et al., 2005; Song et al., 2003) that crosses the blood brain barrier (Alawieh et al., 2018). This intervention partially rescued SVZ proliferation in aged mice, supporting the conclusion that complement-linked signaling contributes to the age-associated reduction in proliferative output. Although insignificant, CR2-Crry treatment reduced microglia inflammation and Mash1+ material within CD68+ compartments in aged microglia. Future work will be aimed at the direct quantification of engulfment dynamics, and mechanistic tests of candidate receptor–ligand axes suggested by the transcriptomic analyses.

Finally, our transcriptomic observations raise a testable hypothesis about target susceptibility. Aging may increase “eat-me” programs in multiple SVZ populations, while “don’t eat-me” programs may be relatively stronger in astrocyte-qNSCs and neuroblasts than in aNSCs and TACs. If validated, this could help explain why increased microglial phagocytic programs in aging are associated with progenitor loss while sparing other SVZ populations. Future work should directly test whether differences in cell-surface inhibitory cues (e.g., *Cd200*, *Pecam1*, MHC class I–linked programs) and corresponding microglial inhibitory receptors shape selective engagement of progenitors in the aged niche, and whether manipulating these interactions can decouple restored proliferation from progenitor loss, potentially paving the way for future therapeutics.

Several limitations should be considered when interpreting these findings. First, our *in vivo* phagocytosis readout relies on confocal colocalization of Mash1 signal within CD68+ compartments in Iba1+ microglia in SVZ wholemounts. While this pattern is consistent with phagolysosomal internalization, additional validation using phagosome markers or ultrastructural approaches will strengthen localization and interpretation. Second, the *in vitro* phagocytosis assay uses pHrodo-conjugated yeast/zymosan-like particles, which can engage receptors such as TLR2 and dectin-1. Consequently, the observed age- and region-dependent differences may reflect pathway-specific responsiveness rather than global phagocytic capacity. Testing alternative cargos (e.g., apoptotic bodies) and defined opsonization states will be necessary to assess generality. Third, although the CR2-Crry intervention is designed to inhibit C3 activation and deposition, we did not directly measure complement activation or target engagement within the SVZ following treatment. This limits causal inference about the extent and spatial specificity of complement inhibition in the niche and makes it difficult to distinguish direct complement effects from downstream or systemic consequences.

Additional caveats apply to generalizability and mechanistic resolution. Our work focuses on discrete ages (3 and 12 months for most experiments), which captures a midlife transition but does not resolve the temporal ordering of microglial activation, phagocytic engagement, and progenitor decline across aging; denser longitudinal sampling would better establish sequence and potential thresholds. The CR2-Crry intervention may not be microglia-specific and could act on other niche cells or peripheral immune pathways, complicating attribution of effects to microglia versus the broader SVZ environment. Moreover, increases in EdU+ counts provide a useful proliferative readout but do not by themselves resolve whether the rescued proliferation reflects altered cell-cycle dynamics, survival, lineage progression, or changes in the composition of proliferating populations; additional phenotyping of EdU+ cells would sharpen interpretation. Finally, the scRNA-seq reanalysis provides supportive mechanistic context, but transcriptomic signatures cannot distinguish increased phagocytic “capacity” from compensatory programs or assign directionality of cell–cell interactions; functional perturbations of candidate receptor–ligand axes and direct measures of engulfment dynamics will be required to establish causality.

## Acknowledgements

This work was supported by funding from the NIH under grants R01NS102448 and T32 AG0201890

## Author contributions

E.K. and R.C. designed the research; R.C., S.H., and N.S.K performed the research; S.T. contributed unpublished reagents/analytic tools; R.C. and S.H. analyzed data; R.C. and E.K. wrote the paper.

