## Supplemental Figures for "Age Associated Increase in Microglia Inflammation and Phagocytosis in the Adult Neural Stem Cell Niche"

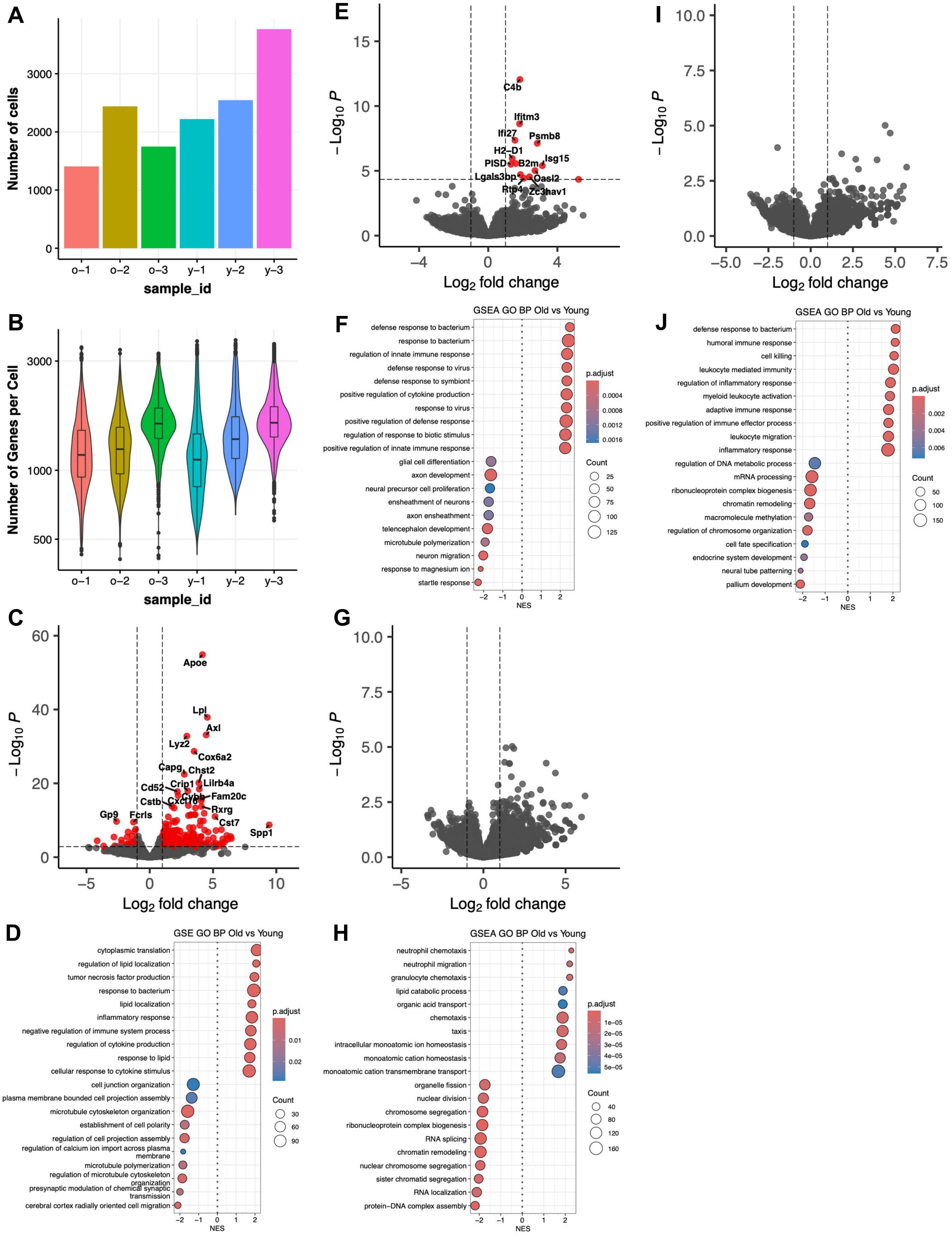


**Supplementary Figure 1. Quality control, differential expression, and gene set enrichment analysis of major cell types in Dulken *et al*., 2019 aging SVZ scRNA dataset**

A) Number of cells captured per sample. O=old (28-29 months old) and Y=young (3 months old), number indicates mouse replicate. B) Distribution of genes identified per cell in each sample. C) SVZ microglia pseudobulk differential gene expression of old vs. young. Red points are $\log_2$ fold change > 1 and adjusted p-value < 0.05. Statistics: n=3 mice/group, DGE results obtained using Wald test followed by Benjamini-Hochberg correction for multiple hypothesis testing. D) SVZ microglia gene set enrichment analysis of gene ontology biological process for old vs. young differentially expressed genes shown in (C). Statistics: GSEA p-values obtained using a permutation test followed by followed by Benjamini-Hochberg correction for multiple hypothesis testing. E-F) Same as in B-C, but for astrocytes-quiescent neural stem cells (qNSCs). G-H) Same as in B-C, but for activate neural stem cells (aNSCs) – neural progenitor cells (NPCs). I-J) Same as in B-C, but for neuroblasts.

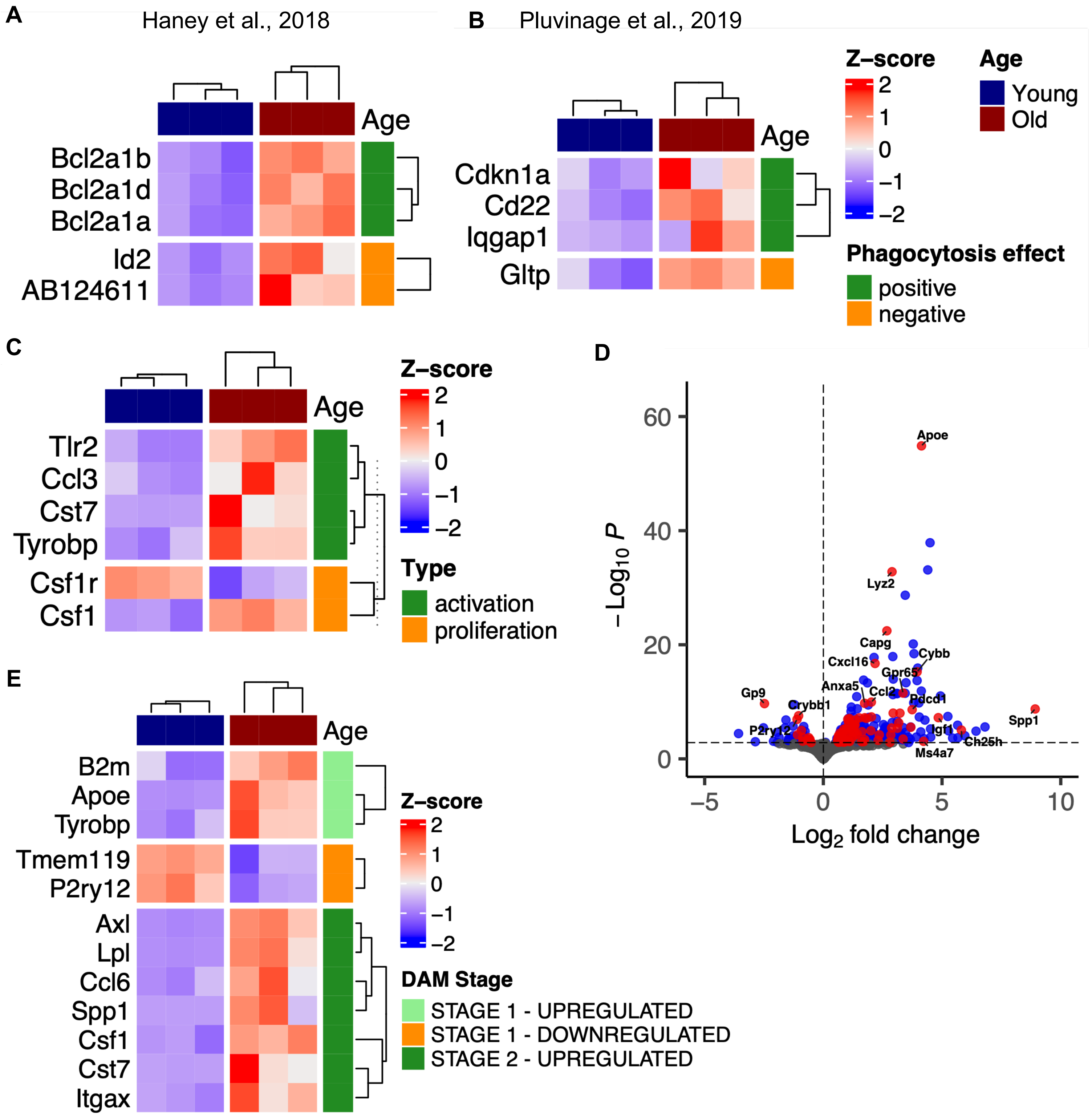


**Supplementary Figure 2. Extended analysis of aging SVZ microglia gene expression changes**

A-B) Heatmap of age-related differentially expressed genes (ARDEGs; adjusted p-value < 0.05) in aging SVZ microglia that overlapped with gene sets derived from screens to identify genes regulating phagocytosis from Haney *et al*., 2018 (A) and Pluvinage *et al*., 2019 (B). Statistics: n=3 mice/group C) Same as in (A), but for ARDEGs that overlapped with the Molecular Signatures Database MICROGLIAL CELL ACTIVATION or MICROGLIAL CELL PROLIFERATION gene sets (Supplementary Table 2). D) SVZ microglia pseudobulk differential gene expression of old vs. young where points highlighted in red are genes that are $\log_2$ fold change > 1 and adjusted p-value < 0.05 and that overlapped with the Molecular Signatures Database entries: DESCARTES FETAL CEREBELLUM MICROGLIA, DESCARTES FETAL CEREBRUM MICROGLIA, DESCARTES MAIN FETAL MICROGLIA, FAN EMBRYONIC CTX BIG GROUPS MICROGLIA, FAN EMBRYONIC CTX MICROGLIA 1,FAN EMBRYONIC CTX MICROGLIA 2, FAN EMBRYONIC CTX MICROGLIA 3 gene sets (Supplementary Table 2). Statistics: n=3 mice/group. E) Same as in (A), but for ARDEGs that overlapped with the disease associated microglia (DAM) gene sets reviewed in Deczkowska *et al*., 2018 (Supplementary Table 2).

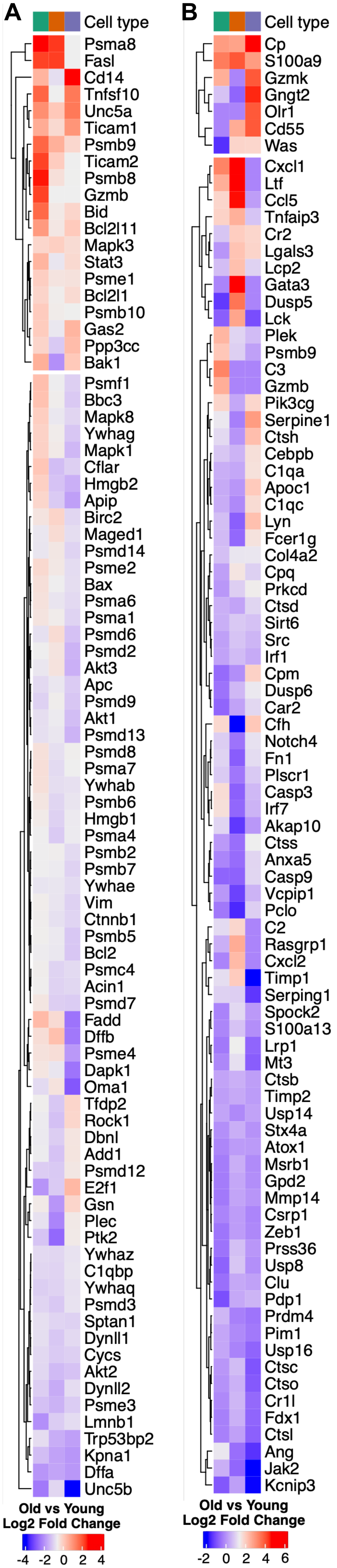


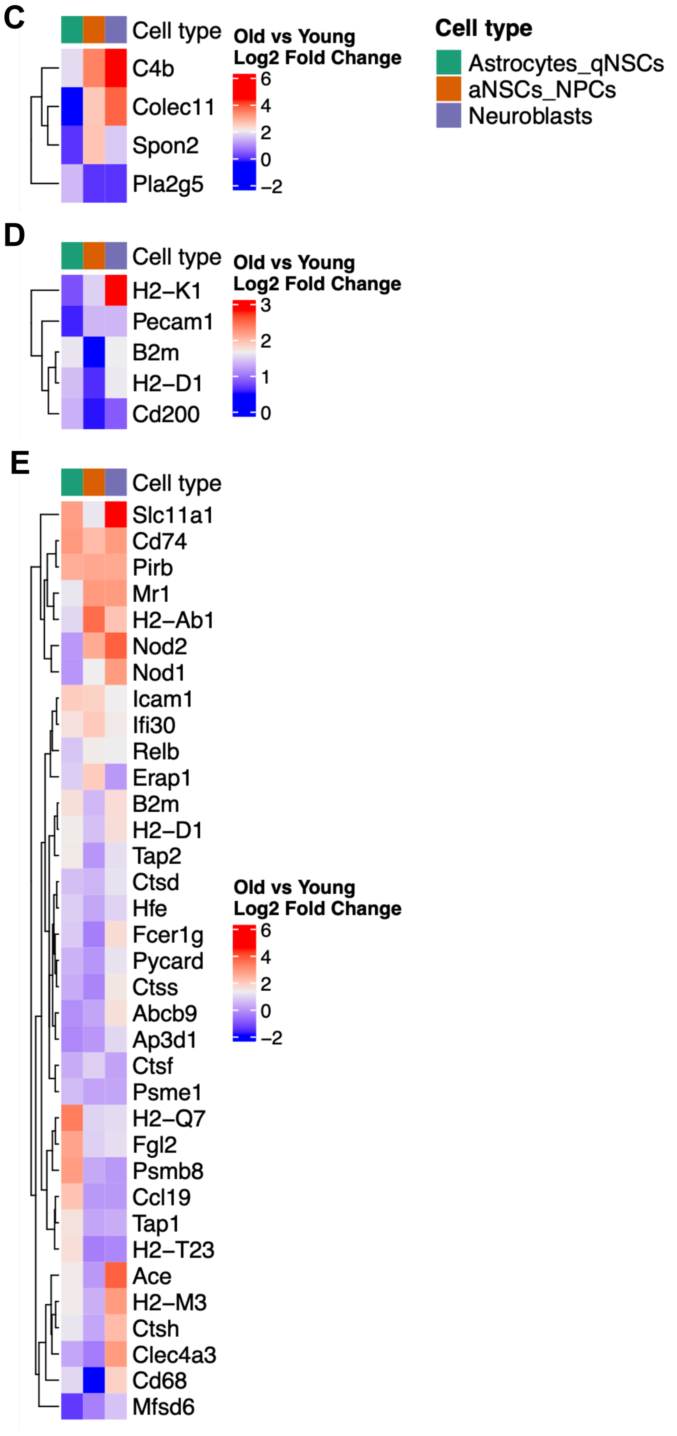


**Supplementary Figure 3. Extended analysis of aging SVZ neural progenitor cell gene expression changes**

A) Average $Log_2$ fold changes in gene expression changes for astrocytes-qNSC, aNSCs-NPCs, and neuroblasts for genes that overlapped with the Reactome Apoptosis gene set (Supplementary Table 2). Fold changes are averages calculated form pseudobulk differential expression analysis (Methods). Statistics: n=3 mice/group. B) Same as in (A), but for genes that overlapped with the Molecular Signatures Database HALLMARK COMPLEMENT gene set (Supplementary Table 2). C) Same as in (A), but for genes that overlapped with the Molecular Signatures Database GOBP OPSONIZATION gene set (Supplementary Table 2). D) Same as in (A), but for genes that overlapped with the curated DON'T EAT ME gene set (Supplementary Table 2). E) Same as in (A), but for genes that overlapped with the Molecular Signatures Database GOBP ANTIGEN PROCESSING AND PRESENTATION gene set (Supplementary Table 2).

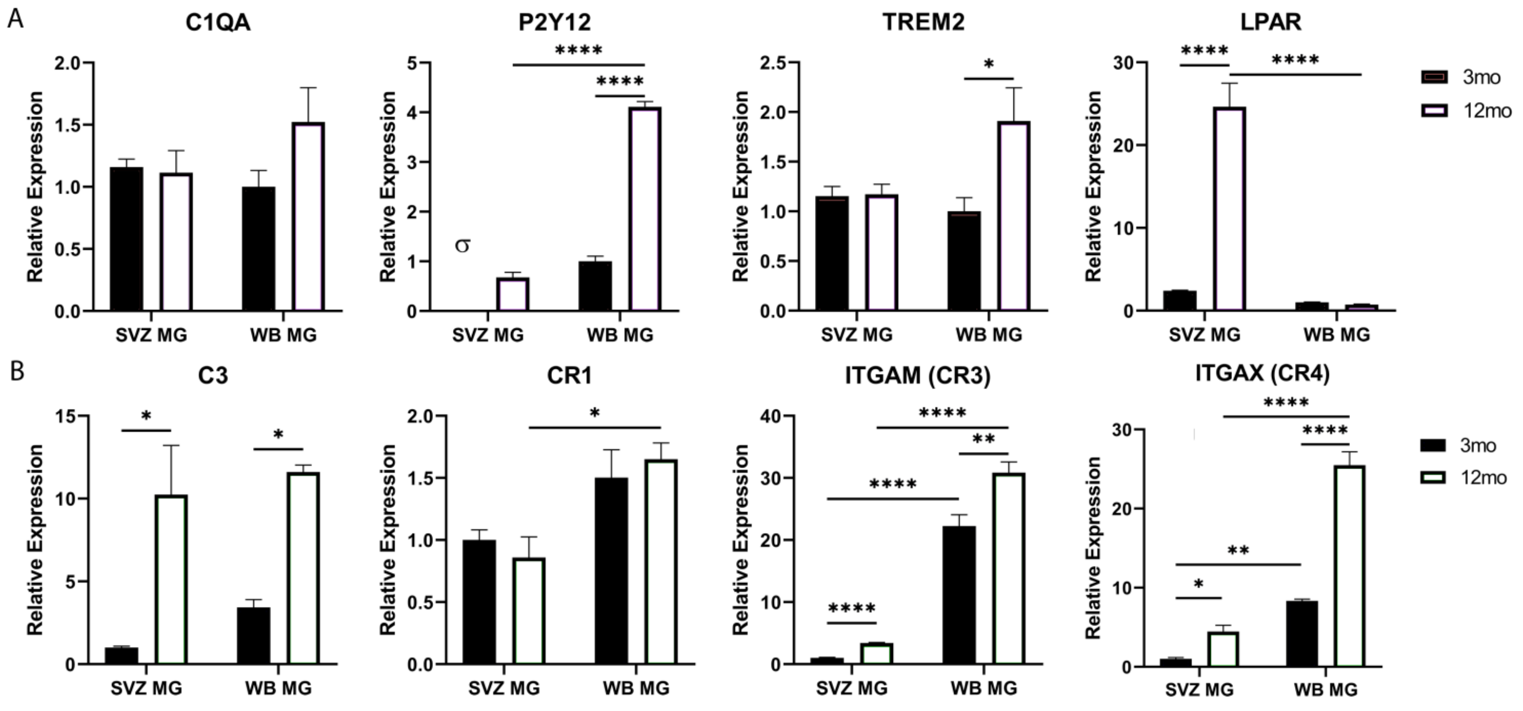


**Supplementary Figure 4. qPCR validation of genes selected from the scRNA-Seq results**
(A-B) Quantification of gene expression of microglia from the SVZ (SVZ MG) or whole brain (WB MG) in young (3 months) and aged (12 months) mice. Statistics: n=3 mice/group; σ: not detected. P-value legend: ns: no symbol; *: p ≤ 0.05, **: p ≤ 0.01, ***: p ≤ 0.001, ****: p ≤ 0.0001.

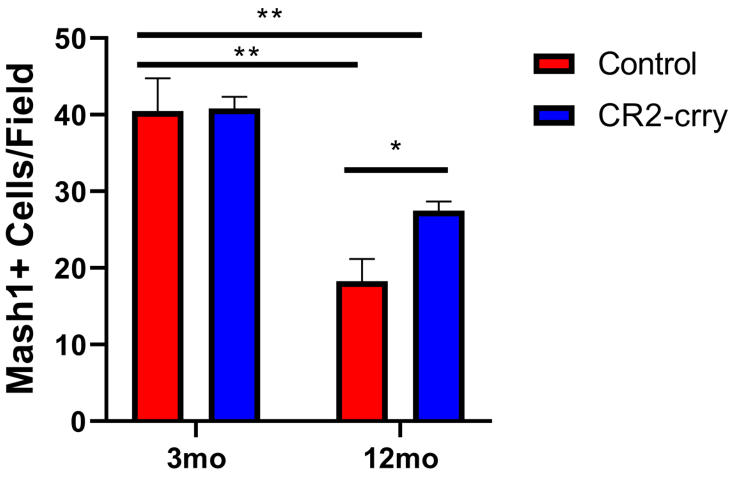


**Supplementary Figure 5. CR2-crry treatment partially rescues Mash1+ cell number**
(A-B) Quantification of proliferating Mash1+ cells in the SVZ of young (3 months) and aged (12 months) mice in response to CR2-cry treatment. Statistics: *n*=4 mice/group, ≥ 100 image fields quantified per mouse, p-values estimated using two-way ANOVA. P-value legend: ns: no symbol; *: $p \leq 0.05$, **: $p \leq 0.01$
