## Supplemental Figure Legends for "Age Associated Increase in Microglia Inflammation and Phagocytosis in the Adult Neural Stem Cell Niche"

**Supplementary Table Legends**

**Supplementary Table 1.** Results of differential expression testing and gene ontology biological process gene set enrichment of old vs. young microglia pseudobulk samples.

**Supplementary Table 2.** Gene sets derived from MSigDB or literature used for analysis of microglia phagocytosis and cells targeted for phagocytosis.

**Supplementary Table 3.** Differential expression results and annotation of genes from the microglia phagocytosis gene set (**Supplementary Table 2**) which were found to be overlap with genes found to be significantly differentially expressed between old vs. young in microglia (**Supplementary Table 1**).

**Supplementary Table 4.** Results of differential expression testing and gene ontology biological process gene set enrichment of old vs. young Astrocyte-qSNC pseudobulk samples.

**Supplementary Table 5.** Results of differential expression testing and gene ontology biological process gene set enrichment of old vs. young aNSC-NPCs pseudobulk samples.

**Supplementary Table 6.** Results of differential expression testing and gene ontology biological process gene set enrichment of old vs. young Neuroblast pseudobulk samples.
